# DLAB - Deep learning methods for structure-based virtual screening of antibodies

**DOI:** 10.1101/2021.02.12.430941

**Authors:** Constantin Schneider, Andrew Buchanan, Bruck Taddese, Charlotte M. Deane

## Abstract

Antibodies are one of the most important classes of pharmaceuticals, with over 80 approved molecules currently in use against a wide variety of diseases. The drug discovery process for antibody therapeutic candidates however is time-and cost-intensive and heavily reliant on *in-vivo* and *in-vitro* high throughput screens. Here, we introduce a framework for structure-based deep learning for antibodies (DLAB) which can virtually screen putative binding antibodies against antigen targets of interest. DLAB is built to be able to predict antibody-antigen binding for antigens with no known antibody binders.

We demonstrate that DLAB can be used both to improve antibody-antigen docking and structure-based virtual screening of antibody drug candidates. DLAB enables improved pose ranking for antibody docking experiments as well as selection of antibody-antigen pairings for which accurate poses are generated and correctly ranked. We also show that DLAB can identify binding antibodies against specific antigens in a case study. Our results demonstrate the promise of deep learning methods for structure-based virtual screening of antibodies.

## Introduction

Antibodies are the most successful class of biotherapeutics, with 85 monoclonal antibody drugs on the US market at the time of writing. Global sales in monoclonal antibody therapeutics reached an estimated $98 billion in 2018 [1], making them one of the fastest growing and largest segments of the pharmaceutical industry. The potential and use of antibodies as therapeutics for a wide range of diseases is due to the high specificity and affinity of their binding, facilitated through the variability in their complementarity determining regions (CDRs) [2].

The development process for novel antibody therapeutics already benefits from computational tools predicting properties of the antibody and the antibody-antigen complex [3]. Here, we demonstrate that machine learning approaches using structural antibody data can enable large scale computational screening in the antibody development pipeline.

In order to develop a successful therapeutic antibody, several features have to be optimised. Antibodies need to bind with high efficacy, specificity and affinity to the target of interest, while at the same time avoiding an immune reaction in the patient [4] as well as avoiding properties that lead to poor developability, such as self-association, viscosity or immunogenicity [5]. To achieve these goals, large-scale experimental screens are usually employed in the pre-clinical stages of antibody-drug development [4, 6].

Typically, initial leads for therapeutic human antibodies are generated using either in vitro display platforms or in vivo transgenic animals.

Further improvement of these initial hits is then achieved through affinity maturation, either via generation and screening of further, hit-based mutagenesis libraries or via rational engineering. High-affinity antibodies generated in this way can be further engineered to achieve desirable properties for antibody therapeutics, for example changes to the constant region to modulate the effector functions as well as *in-vivo* half-life and the improvement of developability [7].

These experimental methods are often effective at generating high-affinity antibodies for downstream development, but are cost-and time-intensive. Furthermore, they do not as standard generate insights into the binding mode of the generated antibodies, or if they are binding to the target epitope on the antigen.

The development pipeline detailed above can be supplemented using in-silico methods, particularly once initial binding candidates have been identified. Computational tools have been used to rationally engineer the binding site in several different ways. For a recent comprehensive review of computational tools employed in antibody engineering, see [8]. As antibody binding affinity is defined by the three-dimensional structure of the antigen-binding region on the antibody (the paratope, which is mainly comprised of the CDRs), antibody modelling tools can aid the design process by rapidly generating models of antibodies (e.g. ABodyBuilder [9], Rosetta [10] and Kotai Antibody Builder [11]). Some methods combine antibody model generation with docking against target antigens and engineering of the antibody (e.g. the Rosetta antibody design tool [12]).

The antibody models generated by those methods, though commonly high quality, are often not accurate across the CDRs, particularly the CDR H3 [9, 13]. This presents a challenge for rigid-body docking methods, which do not allow flexibility of the binding partners, to be able to recapitulate the true antibody-antigen interface. More accurate interface models can be generated using computationally intensive docking approaches [10], however those are too slow to be used on high-throughout screens, where thousands of interface models need to be generated [3].

The caveats described above (CDR model inaccuracy and the speed/accuracy trade-off for interface prediction) mean that currently, no effective structure driven computational tools are available for the early high-throughput screening stages of the antibody development pipeline.

Machine learning approaches in the field of antibody therapeutics discovery have so far mainly focused on sequence rather than on structure as input [14–17]. These machine learning approaches have been shown to be highly efficient for prediction tasks which depend only on the antibody (rather than on the antibody-antigen interface) or are specific to one antigen: random forests have been used both for paratope prediction [14] and for the prediction of VH-VL interface angles in antibody modelling [17]. A current state-of-the art paratope prediction tool, Parapred, is based upon a recurrent neural network approach using the variable region amino acid sequence of the antibody as input data [15]. Another recent study has highlighted the potential of large scale, high throughput machine learning approaches for early stage antibody therapeutics development [16]. A large scale (50k samples) mutagenesis study on one antibody target was used to train a recurrent neural network, which was then used to retrieve new antibody sequences from a sub-sample of antibody sequence space, all of which showed binding affinity *in vitro*. For a review of deep learning approaches in antibody research see [18].

Sequence-based machine learning approaches have yet to provide generalisable predictions across different antigens in one model. An approach in this direction has recently been explored in a study by Akbar *et al*. [19], in which machine-learning accessible descriptions of the binding interface were generated using structure-derived interaction motifs. Further, graph convolutional neural networks have recently been used to encode structural information for the prediction of antibody and antigen interface residues [20], demonstrating the ability of machine learning approaches to successfully utilise structural information derived from both the antibody and the antigen.

Here, we describe a structure-based deep learning approach for early stage virtual screening of antibody therapeutics, when an epitope target of interest is known but no viable hit antibodies have yet been identified. Our approach is able to make generalisable predictions across different antigens. Adopting a three-dimensional gridding method which has been used successfully alongside convolutional neural network methods to predict small molecule/protein binding [21, 22], we implement a similar convolutional neural network trained on rapidly generated rigid-body docking poses of modelled antibody structures in complex with antigen epitopes. We use this **D**eep **L**earning approach for **A**nti**B**ody screening (**DLAB**) to both improve the ranking of docks from the ZDock docking algorithm [23] and, in combination with docking scores generated by ZDock, for the prediction of antibody-antigen binding.

The DLAB source code and pre-trained models are available at https://github.com/oxpig/dlab-public.

## Methods

### Crystal structure data set

The structural antibody database (SAbDab) [24] contains an up-to-date collection of all antibody structures deposited in the PDB [25]. We selected a data set of structures of VH-VL paired antibodies in complex with protein or peptide antigens with a resolution of less than 3 Å from a snapshot of the SAbDab, downloaded on 19/12/2018. The data set consisted of 1216 pdb files of antibody-antigen complexes, of which 759 were non-redundant. Here, we considered an antibody to be non-redundant if its CDR sequence (the concatenated sequence across all CDR regions according to IMGT definition) was only present once in the data set. In the following, this data set of 759 complexes is called the crystal structure data set. The PDB accession codes for the crystal structure data set can be found in Supplementary File 1.

### Model data set

All antibody structures in the crystal structure data set were modelled using ABodyBuilder [9]. Only non-identical template structures were used for modelling. All sidechains were modelled using PEARS [26]. This set of modelled antibodies is referred to as the model data set. Model quality was assessed by calculating the Cα root mean square deviation (RMSD) of each modelled antibody to the crystal structure of the antibody, either across the entire Fv region or across the antibody CDRs by aligning the modelled and the crystal structure antibody by their framework regions and calculating RMSD across the CDR Cα atoms. The antibody models generated using this workflow were of comparable quality to previously published studies [9] (see Supplementary Figure 1A and Supplementary Figure 2).

### Binding and non-binding antibody/antigen examples

We considered every antibody-antigen pair in the crystal structure data set as a binding pair. Since antibodies bind with high specificity, we generated non-binding antibody-antigen pairs by randomly sampling 50 non-cognate antibodies per antigen from the crystal structure/model antibody data set. Our definition of non-cognate required the sampled antibodies to share less than 90% of their CDR sequence with the binding antibody.

### Docking pose generation

For both the crystal structure and model data set, docked antibody-antigen pairs were generated using ZDOCK [23]. For each pair, 500 poses (distinct structures of the antibody-antigen complex) were generated. To aid the docking process, the paratope and epitope were identified and residues not belonging to either the paratope or the epitope were excluded from the interaction site using the standard ZDOCK pipeline as described in the ZDOCK documentation.

The antigen epitope was defined by separating the antibody and antigen in the crystal structure and calculating surface exposed residues for each binding partner separately using the PSA algorithm [27]. All atoms belonging to surface residues on the antigen less than 4 Å from a surface exposed residue on the antibody were considered part of the epitope. Further, all atoms belonging to a surface exposed residue on the antigen within 4 Å of the defined epitope were included in the allowed docking interface to model a realistic level of access to the epitope.

The paratope of crystal structure antibodies was defined in the same way, using the interacting residues from the crystal structure.

For modelled antibody structures, the paratope was defined using the IMGT CDR definition, marking the IMGT defined CDRs and two residues to either side of each CDR as the paratope.

### Clustering for train-test splits

To avoid similarity between binding modes in the train and test sets, CD-Hit [28] was used to cluster antibodies by CDR sequence identity as defined above, using a clustering cutoff of 90% sequence identity. For all learning tasks set out below, train-test splits were performed using clustered cross-validation, assigning all members of a cluster to either the train or the test set.

### Data input for convolutional neural networks

Following the method of Ragoza *et al*. [21], the docking poses were prepared for input into CNNs by discretising the atom information into four-dimensional grids, where three dimensions describe the spatial arrangement of the interaction site and the fourth dimension is used to indicate atom types (see Supplementary Figure 3).

The centre of the interaction site of docking poses was calculated using the PSA algorithm by averaging the coordinates of all surface-exposed atoms within 4Å of the interaction partner on both the antibody and the antigen and taking the mean of the two center points. Poses which after docking had no interactions under 4Å were discarded.

The grid contained only atoms that were within 24Å of the interaction centre. These interaction site specifications were found to cover on average 96% of all interacting atoms on both antibody and antigen (see Supplementary Figure 4). The grid resolution was set to 0.5Å, leading to a total grid size of 96^3^ voxels.

### Reranking ZDOCK docking poses with DLAB-Re

In order to improve ZDOCK docking pose ranking, we created a machine learning method to identify and correctly rank good docking poses. This method, termed DLAB-Re(scoring), is a CNN (architecture shown in Supplementary Figure 5) which predicts the *fnat* score. The *fnat* score is the fraction of contacts between the interaction partners in the crystal structure that are recapitulated in the docked pose. The network generates a probability distribution over 11 *fnat* intervals (10 steps of size 0.1 over [0, 1] and one bin for *fnat* 0.0 poses), which are used to generate a *fnat* prediction via weighted averaging. For model training and testing, the top 500 poses as ranked by ZDOCK for each pairing in the binder set were annotated with their respective *fnat* score interval. For any given antibody-antigen pairing, there are considerably more poses with low *fnat* scores in the top 500 ZDOCK poses than high *fnat* poses. To avoid biasing the network towards predicting all poses into low *fnat* intervals, we used a stratified sampling scheme, sampling poses from each interval at the same rate during training (but not during testing).

During training, the input data was augmented by random rotation around the interaction centre, followed by random translations along the x, y and z axis between −2Å and 2Å. Models were trained for 200,000 parameter update steps using categorical cross-entropy and the rectified Adam optimiser.

Since we wanted to use the improved ranking performance of DLAB-Re on the downstream virtual screening task, it was used to rerank the top 500 poses for all antibody-antigen pairings used during training and evaluation of DLAB-VS. For this experiment, we used the model weights derived during cross-validated training. In the case of cognate and non-cognate antibody-antigen pairings, the DLAB-Re model used was not trained on either the pairings or on the antigen.

To identify antibody-antigen pairings with low quality docking poses, we determined the highest DLAB-Re score given to any of the top 500 poses generated by ZDOCK for each antibody-antigen pairing. This score (DLAB-Re-max) was used to discard particular pairings by ranking all pairings by their DLAB-Re-max score and discarding the bottom 40%, 60% or 80%. To contrast the performance of ZDOCK on the same task, this score thresholding was also applied to the ZDOCK output score of the top pose as ranked by ZDOCK.

### Virtual screening with DLAB-VS and ZDOCK

The goal of virtual antibody screening is to discern binding antibodies against a given epitope from a pool of candidate antibodies. To generate a classification model able to accomplish this task, we trained an ensemble of CNN models (architectures depicted in Supplementary Figure 5), which we termed DLAB-VS (virtual screening), a binder/non-binder classifier for individual docking poses of antibody-antigen complexes.

The input poses for training were selected as follows. For non-binding pairs, the highest ranked pose after DLAB-Re rescoring was selected as a non-binding pose. For binding pairs, we selected up to 50 poses with *fnat* > 0.7 where those were available. Further, following the approach taken by Scantlebury *et al*. [29], 5 poses of the same binding pair with *fnat* < 0.1 were selected as non-binding poses in order to force the networks to learn from the interaction between antibody and antigen by providing for the same antibody-antigen pairing both good and bad binding poses.

Data augmentation was performed in the same manner as for DLAB-Re. Models were trained for 50,000 parameter update steps using the rectified Adam optimiser (as the smaller input data set resulted in earlier convergence). A validation set comprising 10% of the total data set was created using the same CD-Hit clustering as for the training set creation. The validation set was used to select a snapshot of the model during training by choosing the model snapshot with the highest average precision on the validation set. To counteract class imbalance, binder and non-binder poses were sampled so that each batch contained equal numbers of both classes.

For each train/test split and network architecture, two different validation sets were used to train an ensemble of 4 models per fold.

At test time, the DLAB-VS scores of the top 10 poses were averaged. As described above, we used DLAB-Re reranked poses for this purpose. Where an ensemble of models was used, the output scores by the ensemble members were averaged to arrive at the DLAB-VS score. For each antigen target, the antibodies docked against that target (correct and decoys) were ranked by their respective DLAB-VS score.

For the ZDOCK based classifier, the ZDOCK score of the top-ranked binding pose was used to rank the antibodies docked against a particular target.

For the DLAB-VS+ZDOCK model, the DLAB-VS output scores and the ZDOCK output score were normalised per antigen target via minmax scaling and averaged to arrive at the final scores for each target antigen.

The DLAB-Re-max score, output from the reranking method, of each antibody-antigen pairing was used to discard antigens for which the binding antibody was not predicted to have produced any satisfactory docking poses.

### DOVE rescoring

We compared the DLAB-Re results to the DOVE method for CNN-based docking pose ranking [30]. Input file preparation and score generation were performed according to the tutorials on the author’s github page. As detailed on the author’s github page, only the GOAP and ATOM20/ATOM40 scores were used, as the IT-scores were unavailable.

### Additional test sets

To create an unseen test set, which was not used at any point during model choice and hyperparameter optimisation, we used SAbDab entries deposited after the snapshot used for the training data set creation to create an unseen test set. This data set, referred to in the following as the post-snapshot model data set, contained 222 antibody/protein antigen complexes, which formed 173 CDR clusters after clustering the CDR-sequences using CD-Hit at 90% identity. On this test set, we performed modelling, docking, rescoring and binder classification as described above, using the models trained on the model data set, with the exception of the ensemble DLAB-VS score calculation, for which we combined all 40 previously trained models into one ensemble from which the DLAB-VS score for each pairing was averaged. The PDB accession codes for this data set can be found in Supplementary File 2.

We created a SARS-CoV2 data set (the SARS-CoV2 variant data set) by extracting all antibodies from the Coronavirus Antibody Database (CoVAbDab) [31] which were confirmed to bind to the SARS-CoV2 wild-type RBD while also being confirmed not to bind to at least one SARS-CoV2 variant and for which an experimentally determined complex structure was available from which the epitope could be determined as described above. We used ABodyBuilder to model the antibodies as described above. We created structural models of the variant antigens using Foldx5 [32], using the PDB files listed in Supplementary Table 1 as templates and copied the epitope definition for docking purposes from the templates onto the variant models. We then docked each antibody model, defining the paratope as described above, against its epitope on both the wild-type RBD and the confirmed non-cognate variant RBDs and performed rescoring and binder classification as described above, using the 40-model ensemble. As for most of the epitopes in this set, only one antibody was docked, score normalisation was performed over the entire dataset instead of on a per-epitope basis.

### Statistical testing

For statistical significance testing of the difference between means of the best *fnat* in top 10 ranked poses, we used the two-tailed t-test implementation in the scipy python package [33]. For statistical significance testing of the ranking performance of

DLAB-Re, we considered the ratio of antibody-antigen pairs for which a pose with a specific *fnat* is found in the top 10 poses a Poisson rate and calculated p-values using the implementation of the test described in Gu *et al*. [34] in the statsmodels python package [35]. Correspondingly, we approximated the standard deviation of the ratio as 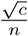, where c is the count of antibody-antigen pairings for which a pose with a specific *fnat* is found in the top X poses and n is the total number of pairings assessed.

## Results

### Crystal structure docking yields high-quality poses

In order to establish a baseline for ZDOCK performance on antibody and antigen crystal structures, we re-docked the complexes in the crystal structure data set and quantified the docking performance through the *fnat* score of the docking poses (Supplementary Figure 1B, C and Supplementary Figure 6). ZDOCK yielded high-quality docking poses, ranking at least one pose with *fnat* > 0.5 in the top ten poses for 93% of the pairings.

### Model docking yields low-quality poses

Docking the model data set antibodies against their cognate antigens on the other hand yields lower quality docking poses. Here, ZDOCK created a pose with *fnat* > 0.5 in the top ten poses for only 44% of antibody-antigen pairings but 70% of pairings had a pose with *fnat* > 0.5 in their 500 highest ranked poses (Supplementary Figure 1C and Supplementary Figure 6).

### DLAB-Re can improve ZDOCK docking pose ranking

Given these results, the first stage was to create a method that is able to identify good docking poses and rank these correctly. DLAB-Re is a CNN trained to predict the *fnat* of docking poses of antibody-antigen pairings. In order to determine the ability of our method, DLAB-Re, to improve docking pose ranking, we ranked the top 500 docking poses generated by ZDOCK for each binder pair by the predicted *fnat* value, using the clustered, cross-validated train-test procedure set out in the Methods section (compare Supplementary Figure 1E).

This rescoring procedure improves upon the performance of ZDOCK ranking. On the crystal structure data set, DLAB-Re recapitulates the ranking performance of native ZDOCK. On the model data set, DLAB-Re increased the number of antibody-antigen pairings for which a pose with *fnat* > 0.5 is ranked in the top ten poses by 16% (p=0.047) (see Fig. 1 C), significantly increasing the mean best *fnat* in the top 10 poses from 0.46 to 0.50 (p=0.002). In Supplementary Figure 7, we show two antibody-antigen pairings for which DLAB-Re strongly increases the *fnat* of the best pose in the top 10 ranked poses.

**Figure 1.**
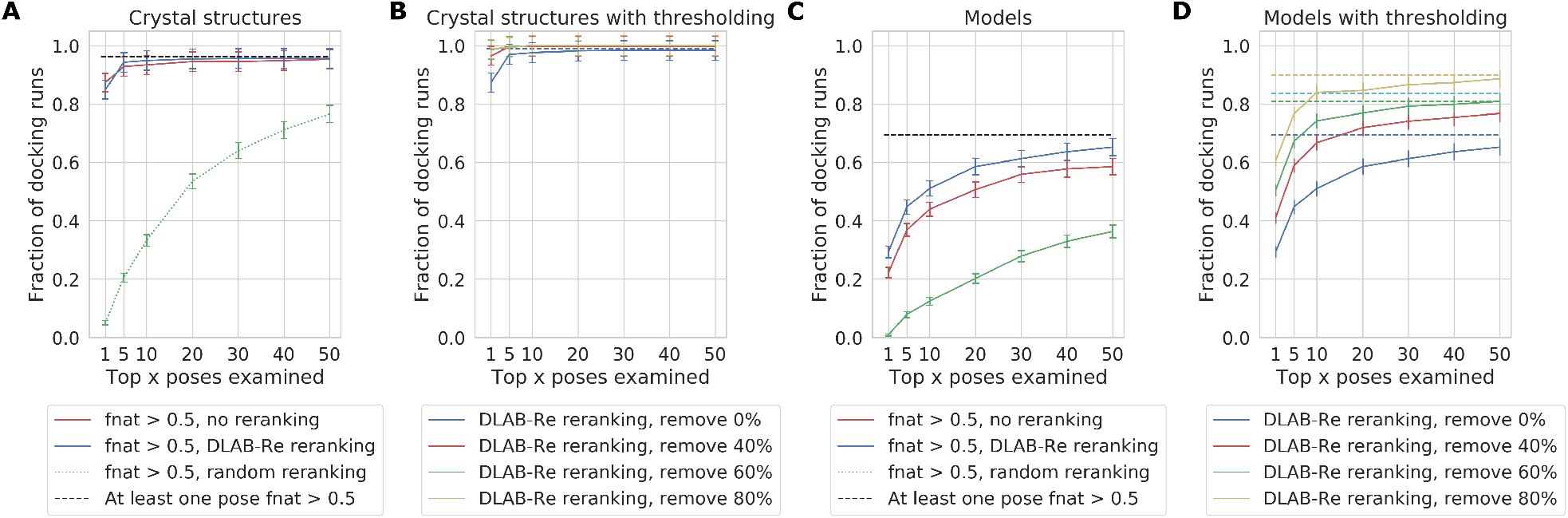
DLAB-Re improves docking performance on the crystal structure data set (**A, B**) and the model data set (**C, D**). On crystal structure data, ZDOCK ranking and DLAB-Re ranking perform similarly and well. On models, the ZDOCK baseline performance is considerably worse and DLAB-Re significantly improves ranking performance. **(A, C)**: DLAB-Re ranks the top 500 poses generated by ZDOCK better than ZDOCK, enriching the ratio of pairings with *fnat* > 0.5 poses ranked highly. The dashed line indicates the fraction of docking runs in which a pose with *fnat* > 0.5 is present in the 500 assessed poses.(**B, D**) Using the DLAB-Re-max score to remove 40%, 60% or 80% of the antibody-antigen pairings respectively can remove antibody-antigen pairings which did not yield high-*fnat* poses. This selects for pairings for which *fnat* > 0.5 poses exist in the top 500 poses generated by ZDOCK (dashed line) and for which DLAB-Re ranks the top 500 poses well (solid line). Error bars are +/- one standard deviation, approximated as described in the Methods section.

### DLAB-Re enables identification of successfully docked antibody-antigen pairings

Using the maximum predicted *fnat* score generated by DLAB-Re, we can discard poorly docked antibody-antigen pairings (see methods and Fig 1B, 1D). Choosing thresholds so that 40%, 60% or 80% of pairings respectively are discarded, the remaining pairings are increasingly enriched both in pairings for which the docking poses are ranked well by DLAB-Re as well as in pairings which have at least one pose with a high *fnat* score in the top 500 poses. Discarding 80% of the pairings in the model data set raises the proportion of pairings for which a pose with at least 0.5 *fnat* was ranked by DLAB-Re in the top ten poses from 51% to 84%, meaning that using this thresholding approach eliminated 93.4% of the antibody-antigen pairings for which DLAB-Re did not manage to rank a pose with at least 0.5 *fnat* in the top ten poses while retaining 33% of the pairings for which it did.

Using this approach with the ZDOCK output scores does not yield the same improvement, only raising the the proportion of pairings for which a pose with at least 0.5 *fnat* was ranked by ZDOCK in the top ten poses from 44% to 57% (see Supplementary Figure 8).

### A CNN docking rescoring tool trained on crystal structure data does not replicate the DLAB-Re performance

We compared the performance of DLAB-Re with the DOVE tool developed by Wang *et al*. [30]. DOVE is a CNN-based docking pose evaluation tool, which is designed to predict docking pose quality according to CAPRI criteria on crystal structure based general protein-protein docking poses. We used the publicly available ATOM40+GOAP model to test whether this training generalises to the antigen -model antibody docking case. Using the DOVE ATOM40+GOAP score to rerank antibody-antigen docking poses, DOVE performs considerably worse than both ZDOCK and DLAB-Re. This holds for both the crystal structure and the model data set (see Supplementary Figure 1D, 9A and 9B).

DOVE is trained to classify docking poses into one of two classes: CAPRI acceptable and not CAPRI acceptable [36, 37], where CAPRI acceptable poses have *fnat* > 0.1 and interface RMSD < 4A or ligand RMSD < 10A. Using this classification to evaluate both DOVE and DLAB-Re results, DOVE still performed worse than ZDOCK and DLAB-Re (see Supplementary Figure 9C). These results highlight the added value from training DLAB-Re both on a domain specific (antibody-antigen) as well as task specific (modelled antibodies) data set.

### ZDOCK easily retrieves binders from the crystal structure data set but not from the model data set

Virtual screening for antibody discovery aims to find binding antibodies for a given epitope from a large set of potential binders. We attempted to retrieve correct binders from the crystal structure data set by docking both the cognate antibody crystal structure as well as 50 non-cognate antibody crystal structures against each antigen crystal structure in the data set. Using ZDOCK, the cognate binder is found in the top 2% (i.e. top ranked) of antibodies for 49.7% of antigen targets, and in the top 10% in 65.6% of antigen targets, outperforming the random baseline. On the model data set, using the same procedure, ZDOCK ranks the binder in the top 2% of antibodies for only 5.5% of antigen targets, and in the top 10% for only 18.8% of antigen targets (see Supplementary Figure 10).

### DLAB-VS and ZDOCK can be combined to improve performance on the model data set

In order to improve our ability to perform virtual screening on modelled antibodies, we trained a new model, termed DLAB-VS (virtual screening) to classify antibody-antigen pairings as binders or non-binders, as detailed in the methods section. For a large proportion of model antibody structures, the docking pipeline does not yield high quality complex structures (see Fig. 1). Therefore, at train time only docking poses with *fnat* > 0.7 were shown to the network as positive binders and poses with *fnat* < 0.1 were shown to the network as non-binders. Furthermore, at test time we averaged the network output over the ten highest ranked poses for each antibody-antigen pairing. Lastly, rather than training a single model for each train/test fold, we trained four models using two different architectures for each of ten clustered cross-validation folds as detailed in the methods section and used the averaged output as the DLAB-VS score. Using this training approach, the DLAB-VS model achieved classification performance comparable to ZDOCK: on the model data set, the cognate antibody was ranked in the top 2% of antibodies for 4.7% of antigens and in the top 10% for 16.4% of antigens (see Supplementary Figure 10).

However, the two approaches, ZDOCK and DLAB-VS, did not perform equally across antigen targets. It was possible to improve classification performance by using the mean of the two scores to rank putative binders (see methods and Supplementary Figure 1E and 1F). Using this approach, termed DLAB-VS+ZDOCK, the binder was ranked in the top 2% of antibodies for 6.4% of antigen targets, and in the top 10% in 19.7% of antigen targets (see Fig. 2). In the following, we use this DLAB-VS+ZDOCK approach.

**Figure 2.**
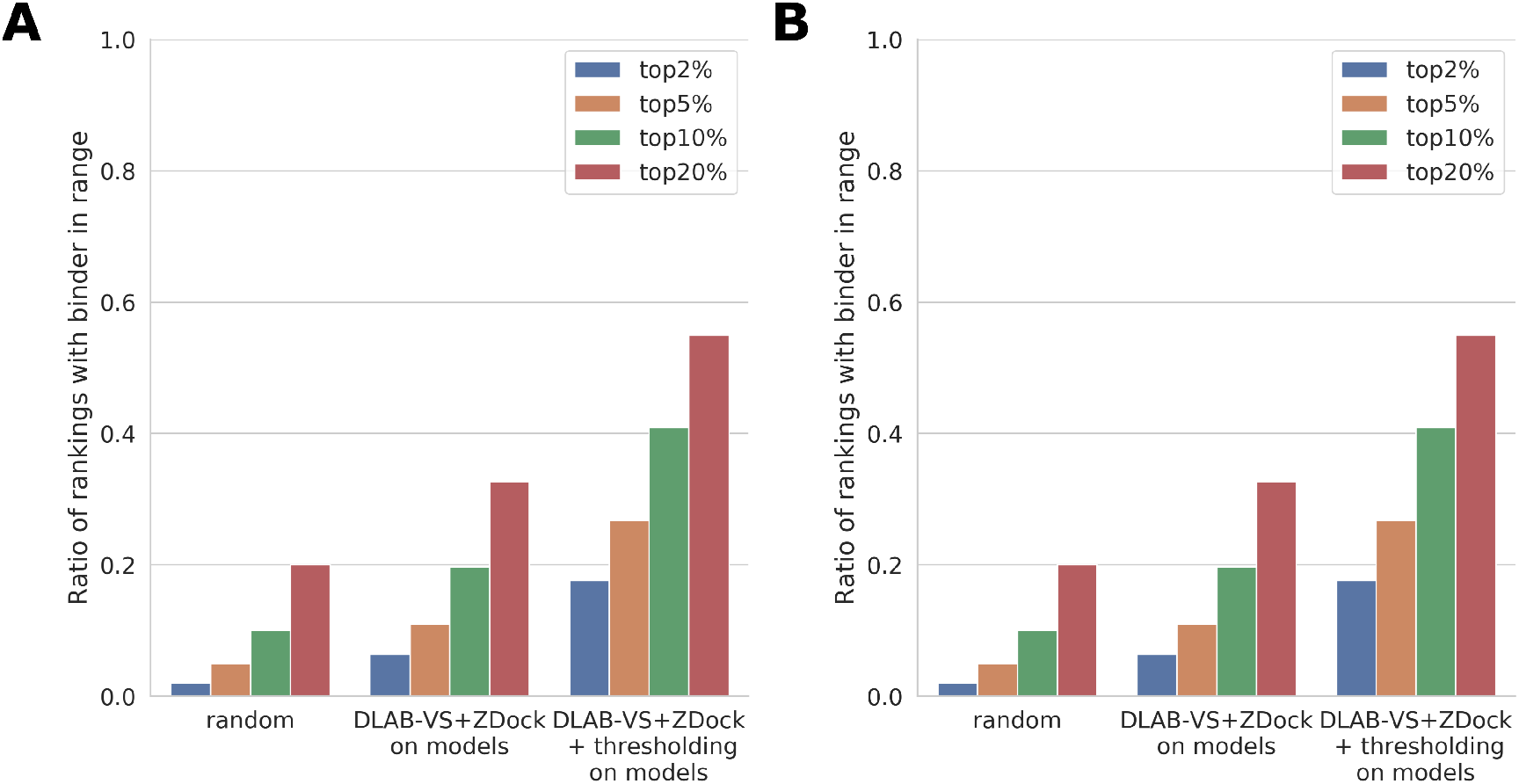
DLAB-VS and ZDOCK binder classification performance. For each approach, the ratio of pairings for which the binding antibody was ranked in the top 2%, top 5%, top 10% and top 20% respectively is shown. **(A)**: Comparison of the performance of ZDOCK and DLAB-VS binder classification on the model data set to the random expectation (“random”) of finding the binder in the top N%. Using the combination of DLAB-VS and ZDOCK scores (“DLAB-VS+ZDOCK”) detailed in the methods section and supplementing it with the DLAB-Re-max thresholding (“DLAB-VS+ZDOCK + thresholding”), the classification performance on the model data set can be improved significantly. **(B)**: Performance on the post-snapshot model data set after removing any CDR sequences with overlap to the model data set (at 90% CDR sequence identity as defined above) both without (“DLAB-VS+ZDOCK”) and with (“DLAB-VS+ZDOCK + thresholding”) DLAB-Re-max score thresholding. The DLAB-VS performance on this test set is better than the performance on the model data set, as discussed in the Results section. Performance for the CDR sequences clustering with sequences in the model data set is shown in Supplementary Figure 12.

### Using DLAB-Re to discard antigen targets enables selection of well-performing models

The performance of the DLAB-VS+ZDOCK model is highly dependent on the quality of the docking poses from which the score is derived as well as the quality of the antibody model (see Supplementary Figure 1G and 11). As described above, DLAB-Re enables the selection of well-docked antibody-antigen pairings, therefore, to further improve the performance of the DLAB-VS+ZDOCK classifier, we used the output from DLAB-Re to identify antibody models with high likelihood of being well docked from the model data set. To test if this would improve results, for each of the training cross-validation folds, we only considered antigen targets for which the DLAB-Re-max score of the cognate antibody was within the top 20% of DLAB-Re-max scores of antibody-antigen pairings within that fold. On these targets, where the cognate antibody was predicted to be well docked by DLAB-Re, the cognate antibody was ranked in the top 2% for 17.6% of antigens and in the top 10% for 40.8% of antigens (see Fig. 2A). This improvement was reliant on using the combined DLAB-VS+ZDOCK model, using the same approach while ranking by the ZDOCK output scores alone only increased the classification performance marginally (see Supplementary Figure 10).

### DLAB-VS+ZDOCK performs well on the post-snapshot model data set

In order to test the performance of the DLAB pipeline on a completely unseen test set (see Supplementary Figure 1H), we ran the pipeline on the post-snapshot model data set (see methods). On this set, an ensemble of all 40 DLAB-Vs models trained during the clustered cross-validation training on the model data set and the ZDOCK output scores were used to rank the cognate antibody model as well as 50 non-cognate antibody models. The ensemble achieved higher performance on this test set than on the previous test cases (both with and without the DLAB-Re selection criterion).

In order to calculate how different the post-snapshot model data set was from the training set and the potential for this to influence model performance, we clustered the antibody CDR sequences in both the model data set and the post-snaphot model data set at 90% identity. Of the new CDR sequences, 17.3% clustered with at least one CDR sequence in the snapshot. On the cognate antigen targets for those antibodies, the large ensemble performs exceptionally well both before and after DLAB-Re thresholding (binder in top 2% for 18% and 57% of antigen targets respectively, see Supplementary Figure 12). On the subset of new additions to SAbDab without overlap to the snapshot, the performance using the 40-model DLAB-VS+ZDOCK ensemble is similar to the performance on the snapshot using 10 folds of 4-model ensembles (see Fig. 2B). The generalisation performance of DLAB-VS is therefore not based on overlap between the training set and the post-snapshot test set. We further demonstrated this by assessing the performance of the DLAB pipeline on the set of post-snapshot targets with cognate antibodies with at most 85%, 80%, 75% or 70% CDR sequence identity to any antibody in the training set. We observed no significant change in performance between the different thresholds, demonstrating that the predictive performance of DLAB is generalisable (see Supplementary Figure 13).

### DLAB-VS can distinguish binding and non-binding antigen variants

A use case of interest is determining whether mutations in the antigen can disrupt antibody binding (see Supplementary Figure 1I). In order to test whether this task is accessible to structure-based deep learning tools, we created a data set of antibodies confirmed to bind against the SARS-CoV2 wild-type receptor binding domain (RBD) while also being confirmed not to bind against at least one SARS-CoV2 RBD variant. We ran the data set through the DLAB pipeline as described in section 3.10 and assessed whether the DLAB-VS+ZDOCK output score consistently scored the antibody-wild-type pair higher than the antibody variant pair. For the 14 antibody-variant pairs in the data set, the DLAB-VS+ZDOCK score of the antibody-wild-type pair was higher than the score of the antibody-variant pair in 13/14 cases. This result indicates that the variant classification problem is accessible to structure-based deep learning tools.

## Discussion

One of the major shortcomings of current computational antibody drug discovery is the lack of structure-based, early-pipeline screening tools to identify promising candidate antibodies.

Here, we have shown how DLAB, our structure-based deep learning approach, can be used to improve pose selection in antibody-antigen docking experiments and can enable the identification of antibody-antigen pairings for which accurate poses have been generated and selected. DLAB-Re is able to identify pairings for which a pose with *fnat* > 0.5 is in the top ten poses for 84% of the pairings, which can be used to improve binder classification performance downstream.

We have furthermore demonstrated that our DLAB tool can identify putative binders to a given epitope in several different settings. The complete DLAB pipeline of docking followed by DLAB-Re and DLAB-VS enriched binders both against the background of non-binding SAbDab-deposited sequences as well as in a more realistic usage scenario against H3 length-matched antibody sequences drawn from antibody repertoire data.

On the crystal data set with highly accurate antibody structures and docking poses, both DLAB-VS and ZDock are able to strongly enrich binders. In the case of model antibodies docked to antigens, where both model and docking quality have to be considered, ZDock and DLAB-VS approaches fail to achieve strong discrimination between binders and non-binders. However combining the two scores improved performance. These results are in line with previously published work on the ability to classify cognate antibodies through cross-docking analysis [38].

On a realistic use case, using the SARS-CoV2 receptor binding domain epitope as the target epitope, we have demonstrated the utility of the DLAB pipeline, correctly scoring antibody escape variants lower than the cognate epitopes for 13 of 14 antibody-variant pairs.

The DLAB pipeline has been trained specifically on a combination of ABodyBuilder and ZDOCK. The use of a different input pipeline would likely require additional finetuning of the weights of both DLAB-Re and DLAB-VS. One natural extension would be the use of flexible docking approaches, which could improve the input docking poses but would be computationally expensive given the scale of experiments needed in a high-throughput setting.

We have demonstrated the applicability of structure-based deep learning approaches both to antibody research in general and to the virtual screening task specifically. Methods such as DLAB will improve with increasing availability of structural antibody data as well as improved antibody modelling and improved fast docking methods. DLAB demonstrates the potential of structure-based deep learning approaches to supplement traditional experimental screening approaches and sets a course for structure-based virtual screening methods for antibody drug discovery.

## Acknowledgements

This work was supported by funding from the Engineering and Physical Sciences Research Council (EPSRC) and the Medical Research Council (MRC) [grant number EP/L016044/1] and AstraZeneca.

## Supplementary Information

**Supplementary Figure 1.**
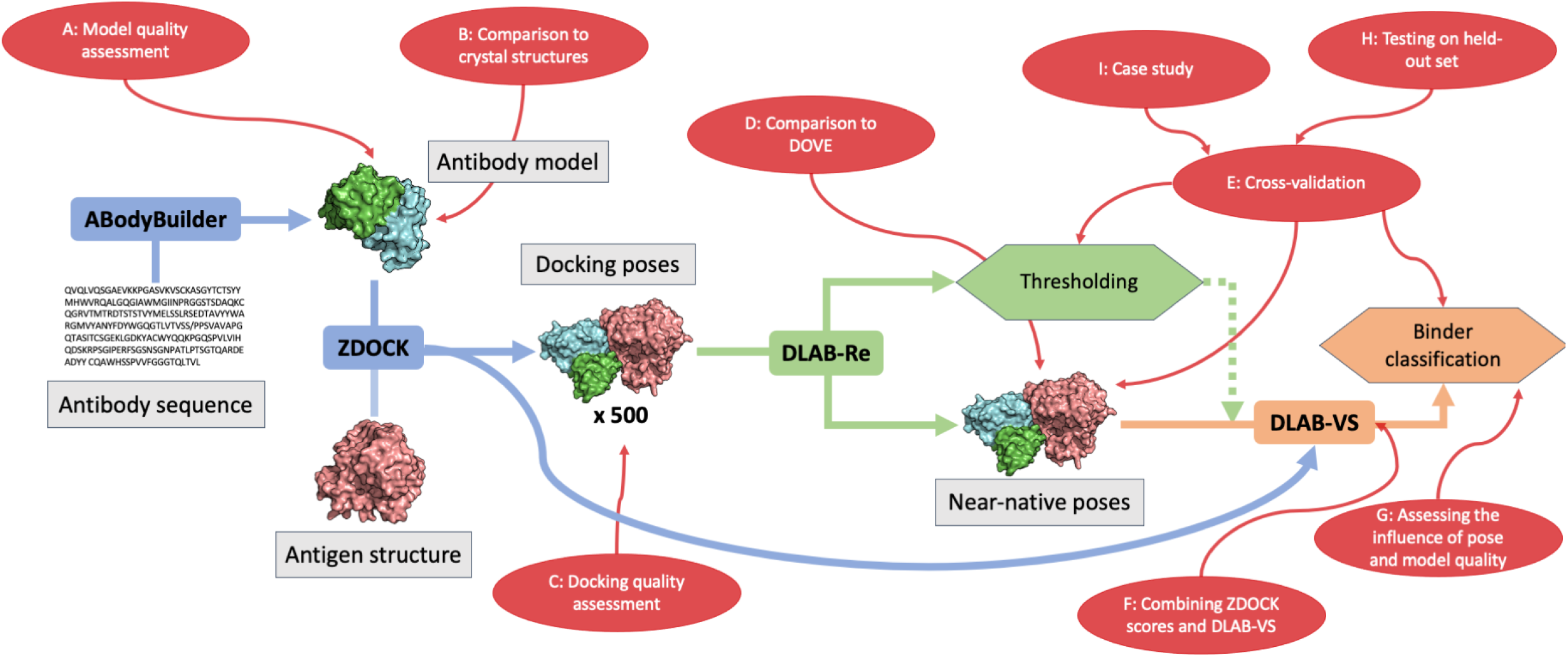
Flowchart depicting the DLAB pipeline and considerations during the development of the final DLAB ensemble model. Blue boxes/arrows indicate external tools, green boxes/arrows indicate where DLAB-Re fits in the pipeline and similarly, orange boxes/arrows indicate DLAB-VS. Red boxes/arrows and the corresponding labels highlight the points of the pipeline analysed during different stages of the model development process and are referred to throughput the main text body.

**Supplementary Figure 2.**
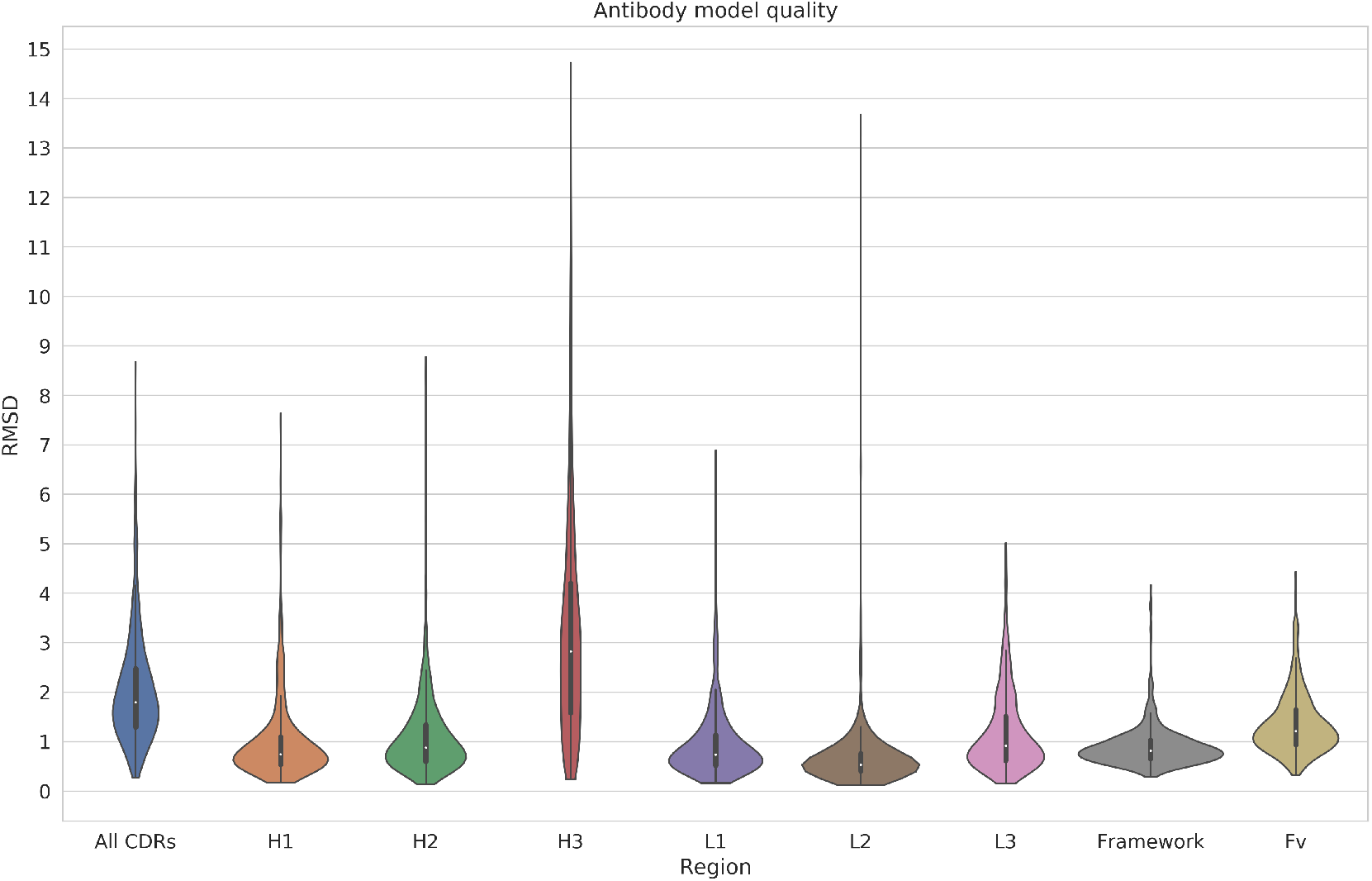
Assessment of the ABodyBuilder generated antibody models. RMSD values across regions of interest on the antibody are calculated as described in the Methods section of the main paper.

**Supplementary Figure 3.**
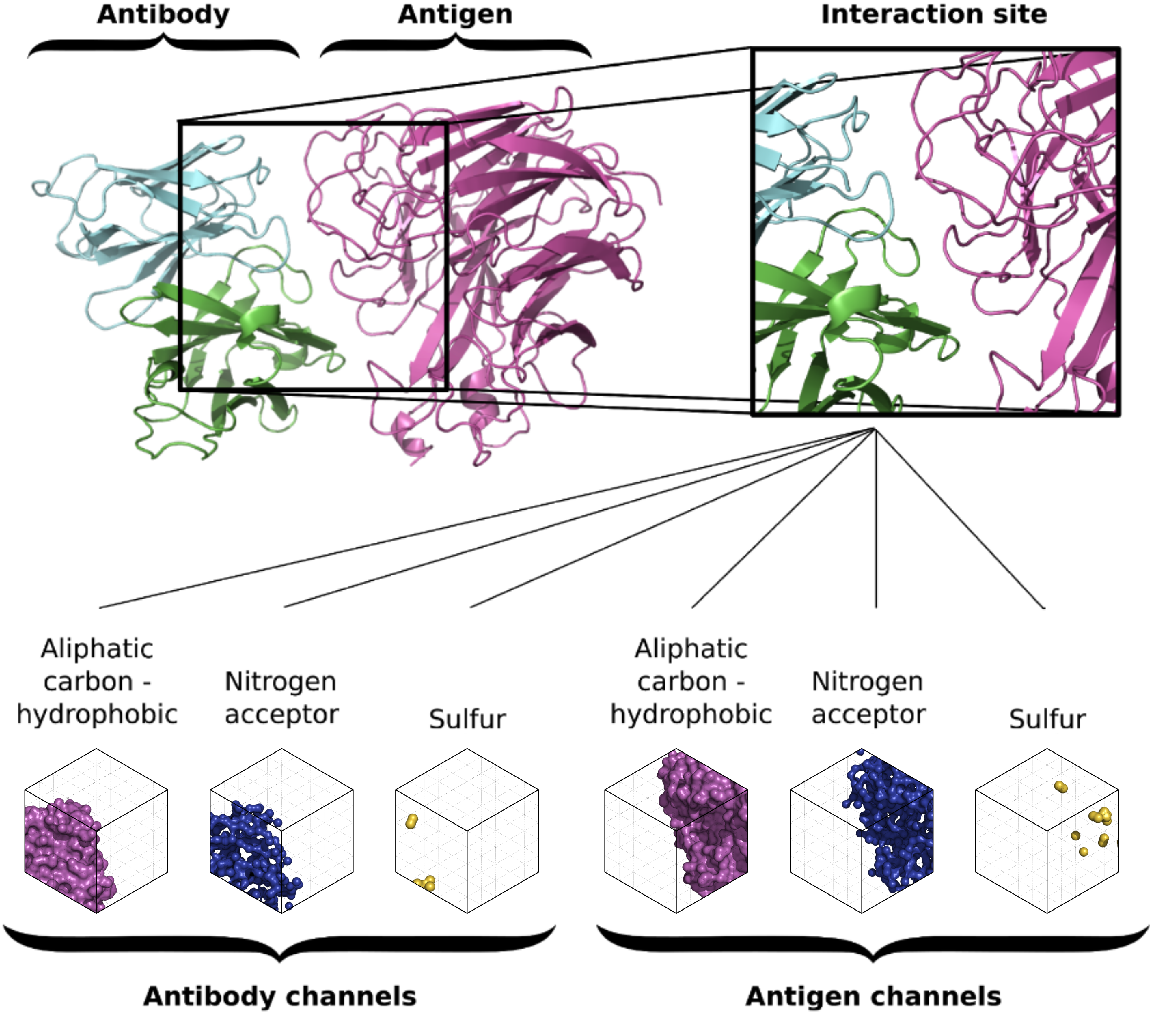
The binding interface discretisation algorithm. (**Top**) The interface is defined as a 48Å cube centered on the interaction center between antibody and antigen. (**Bottom**) The atoms occupying the interface in the docking pose are discretised using the libmolgrid python API into 3D arrays of atom type densities (14 types for both antibody and antigen, of which 3 are depicted here as an example).

**Supplementary Figure 4.**
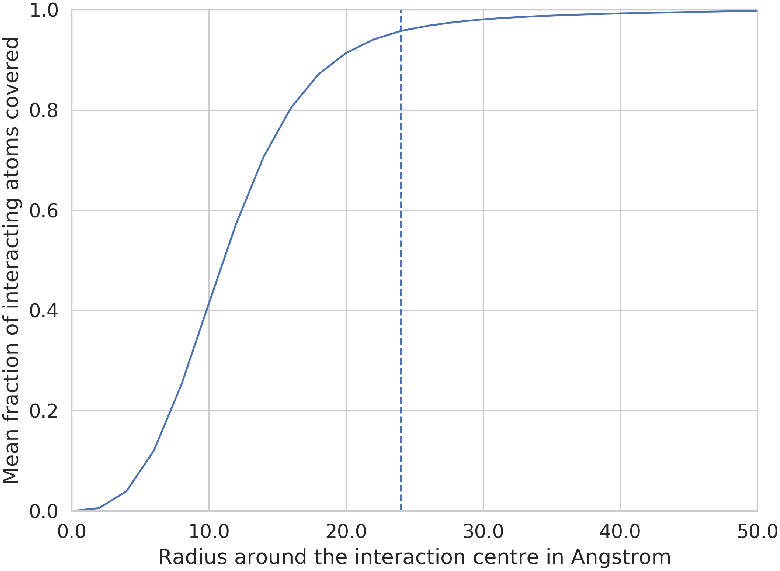
The average fraction of interacting atoms (between the antibody and antigen) included in the interaction box depending on the radius around the interaction center. The dashed vertical line indicates 24Å, the radius used in this study.

**Supplementary Figure 5.**
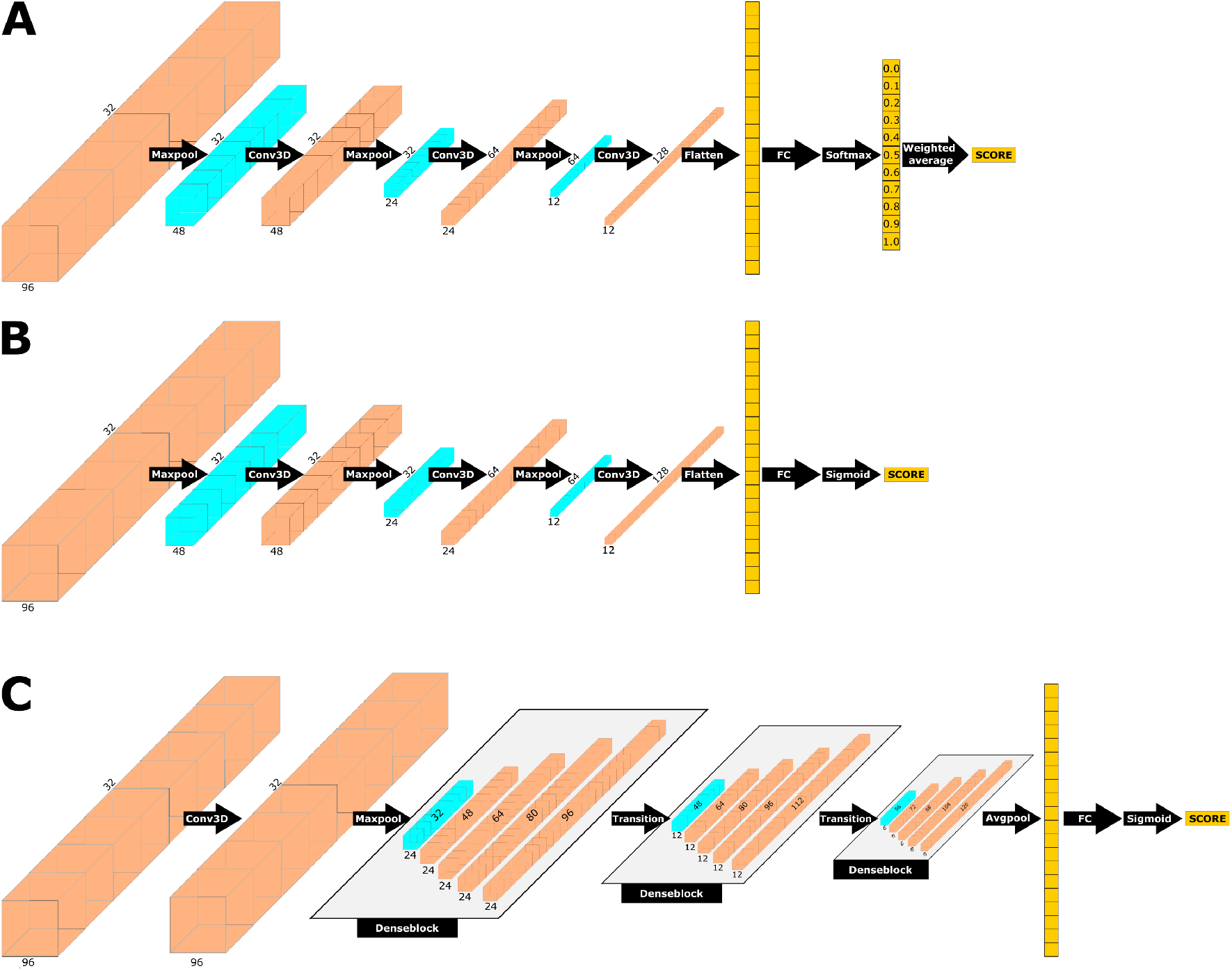
CNN architectures used in the study. (**A**) DLAB-Re architecture. A 3-layer CNN followed by a fully connected layer is used to predict membership of one of 11 *fnat* intervals. To make *fnat* predictions, the output of the softmax layer is transformed into a single score via weighted averaging. (**B, C**) The DLAB-VS architectures forming the ensemble of DLAB-VS models. Both architectures output a binary binder/non-binder classification for an antibody/antigen pairing. (**B**) The same CNN architecture as in (A) with a different output layer. (**C**) A CNN architecture consisting of a single convolutional layer followed by 3 Denseblocks, using the same output layer as (B).

**Supplementary Figure 6.**
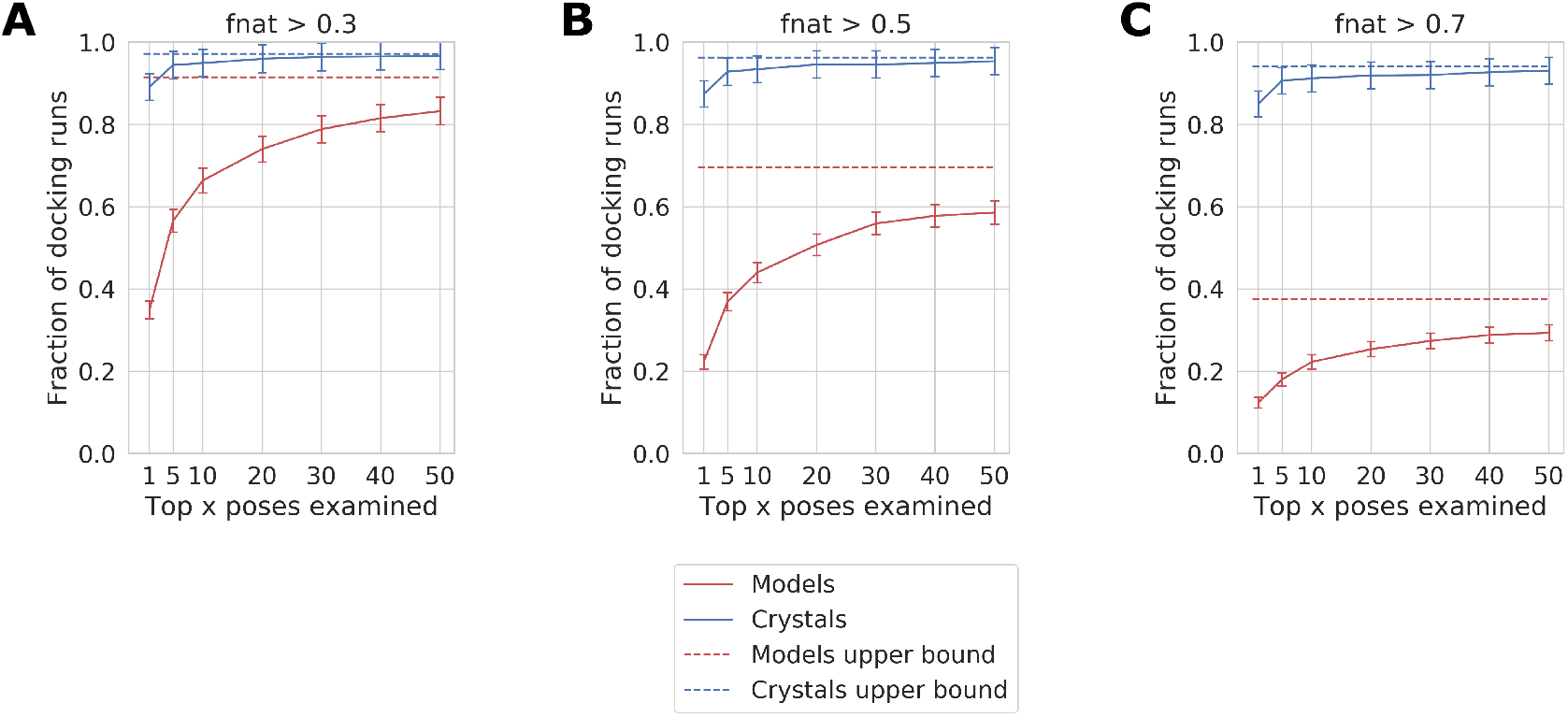
Comparison of ZDock runs on crystal structures and models. Each of the three plots shows the fraction of antibody-antigen pairings which have at least one pose meeting the threshold *fnat* score 0.3 (A), 0.5 (B) or 0.7 (C) in their x highest ranked docks. The dashed line indicates the fraction of docks which have such a pose in their top500 poses (the upper bound of the pose ranking algorithm.

**Supplementary Figure 7.**
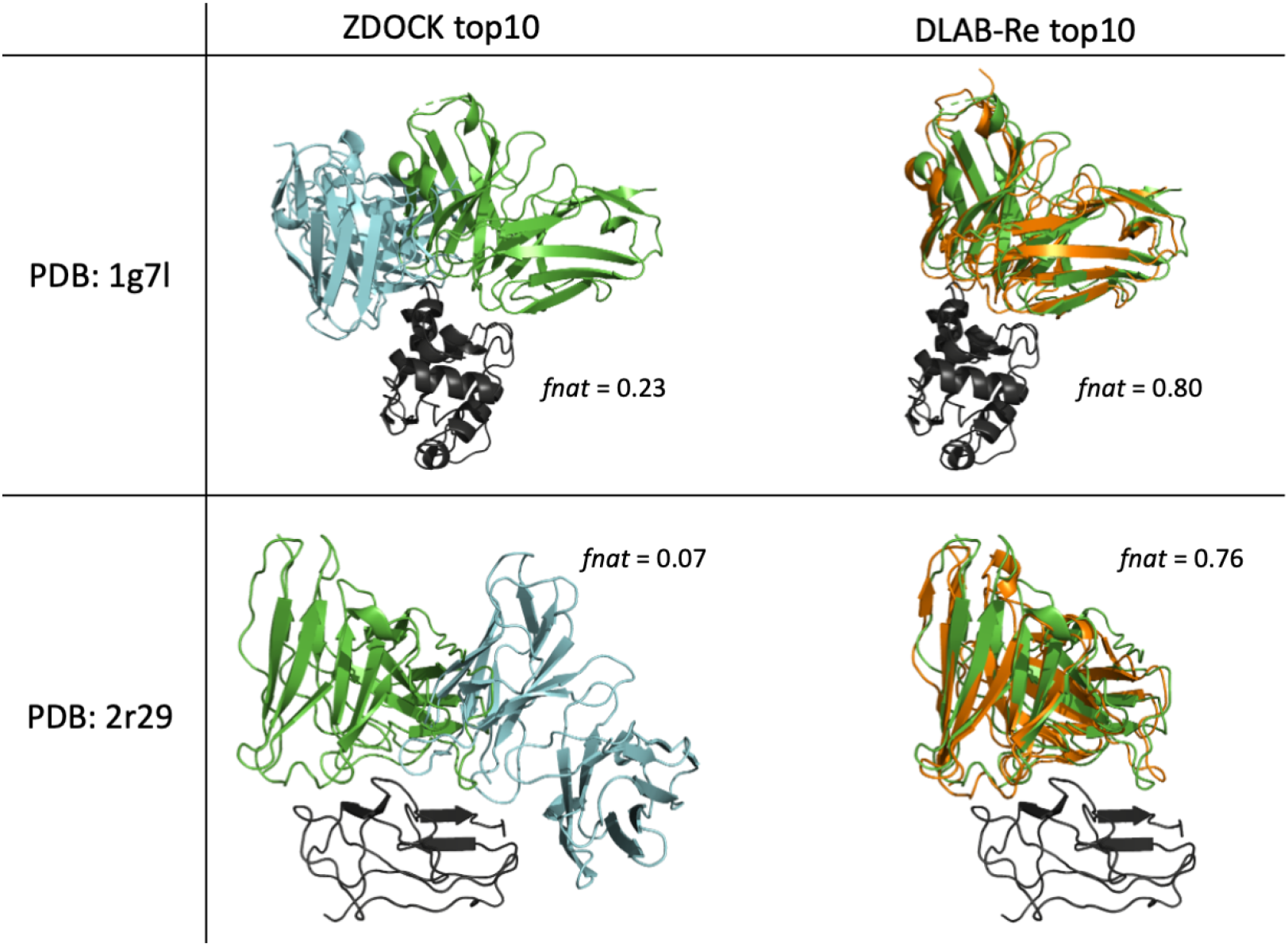
Two examples for which DLAB-Re strongly improves pose ranking for modelled antibodies. (Left) Pose with the highest *fnat* in the top 10 poses as ranked by ZDOCK. (Right) Pose with the highest *fnat* in the top 10 poses as ranked by DLAB-Re. The target antigen is depicted in black, the experimentally determined bound antibody structure in green, the pose with the highest fnat in the top 10 poses as ranked by ZDOCK in cyan and the pose with the highest fnat in the top 10 poses as ranked by DLAB-Re in orange.

**Supplementary Figure 8.**
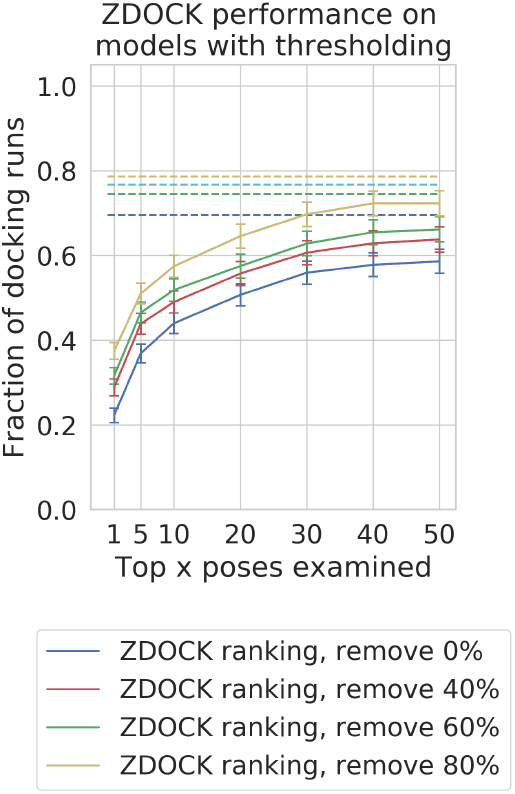
Performance of ZDock-based max score thresholding on ranking performance. While higher overall ZDock scores correlate with improved ranking performance, the effect is much less pronounced than using DLAB-Re-max scores. The solid lines indicate the fraction of antibody-antigen pairings which have at least one pose with *fnat* over 0.5 in their x highest ranked docks, the dotted line indicates the fraction of pairings for which a pose with *fnat* over 0.5 exists in top 500 poses.

**Supplementary Figure 9.**
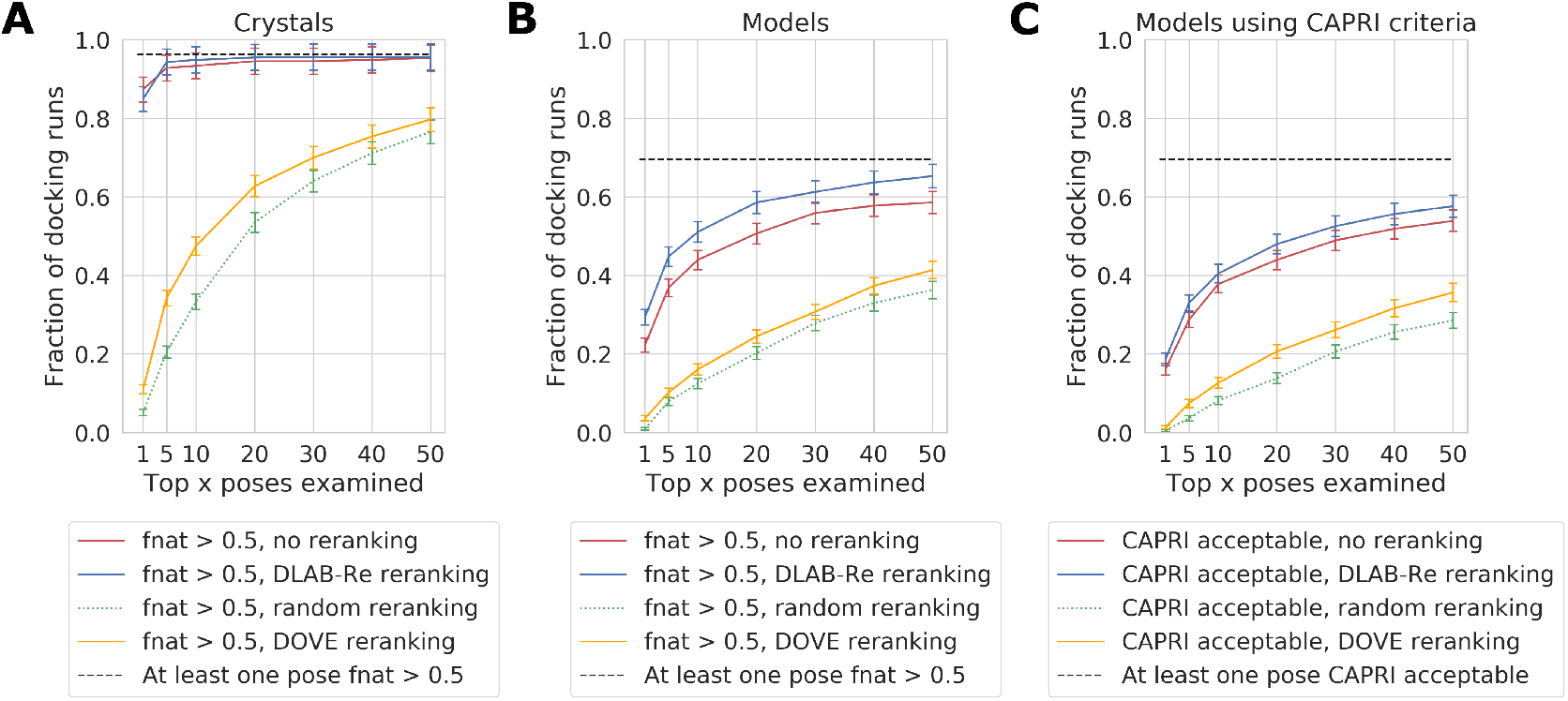
Comparison of DLAB-Re performance to DOVE performance. (A, B) Comparison using the pose *fnat* scores on the crystal structure data set (A) and the model data set (B). The fraction of antibody-antigen pairings which have at least one pose with *fnat* over 0.5 in their x highest ranked docks for ZDock ranking (“no reranking”) as well as DOVE and DLAB-Re reranking is shown in solid lines, the baseline of randomly shuffling the top 500 poses is shown with a dotted green line and the ratio of pairings with a pose with *fnat* over 0.5 is indicated with a dotted black line. (C) Comparison using the CAPRI acceptability criterion for each pose. Legend explanation see (A, B). In all three cases (A, B, C), DLAB-Re achieves an improvement over native ZDock, while DOVE does not replicate the ranking performance achieved by ZDock.

**Supplementary Figure 10.**
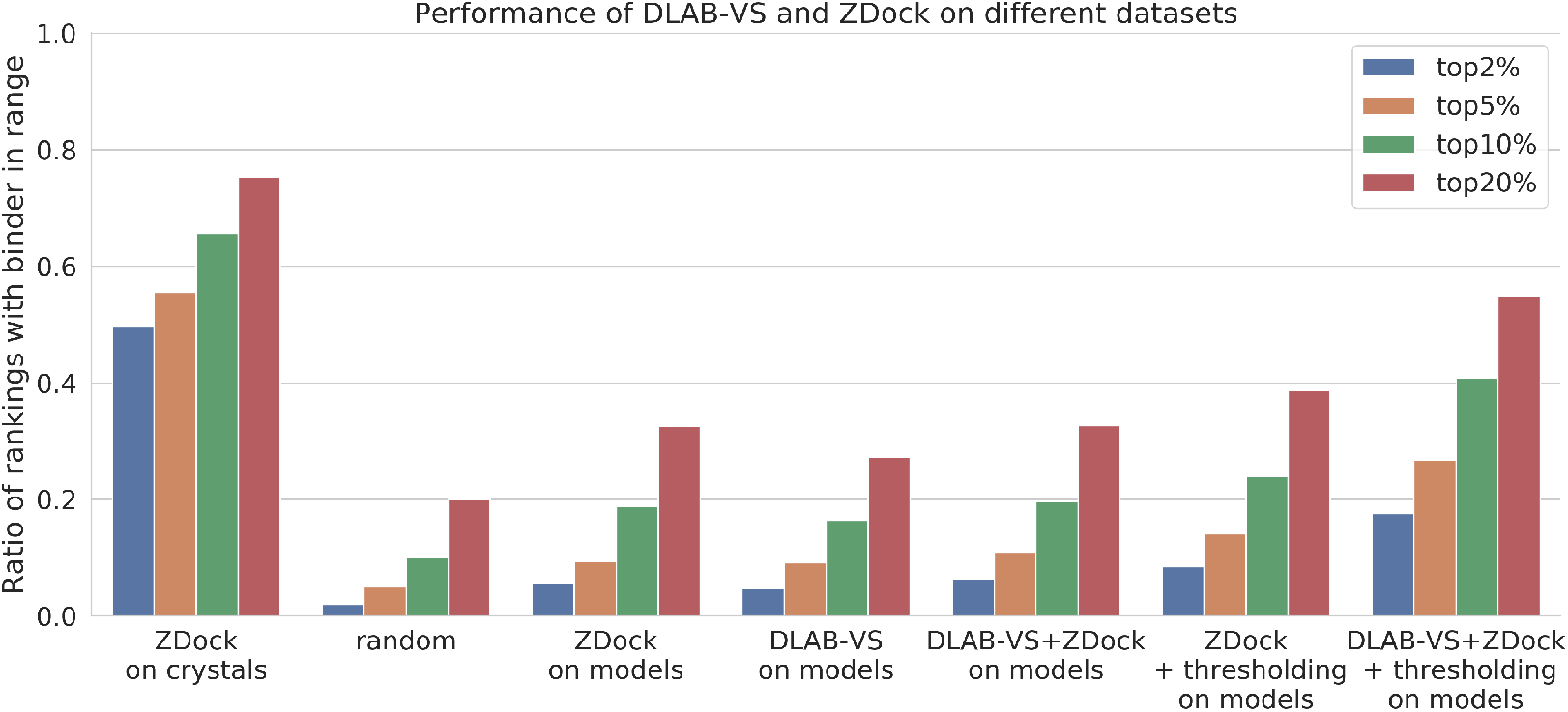
DLAB-VS and ZDock binder classification performance on different data sets. For each approach, the ratio of pairings for which the binding antibody was ranked in the top 2%, top 5%, top 10% and top 20% respectively is shown. Comparison of the performance of ZDock binder classification on the crystal data set to the random expectation (“random”) of finding the binder in the top N% and the performance of ZDock, DLAB-VS and the combination of ZDock and DLAB-VS generated as detailed in the Methods section (“DLAB-VS+Zdock”) on the model data set, both with and without using the DLAB-Re-max thresholding approach (“+ thresholding”).

**Supplementary Figure 11.**
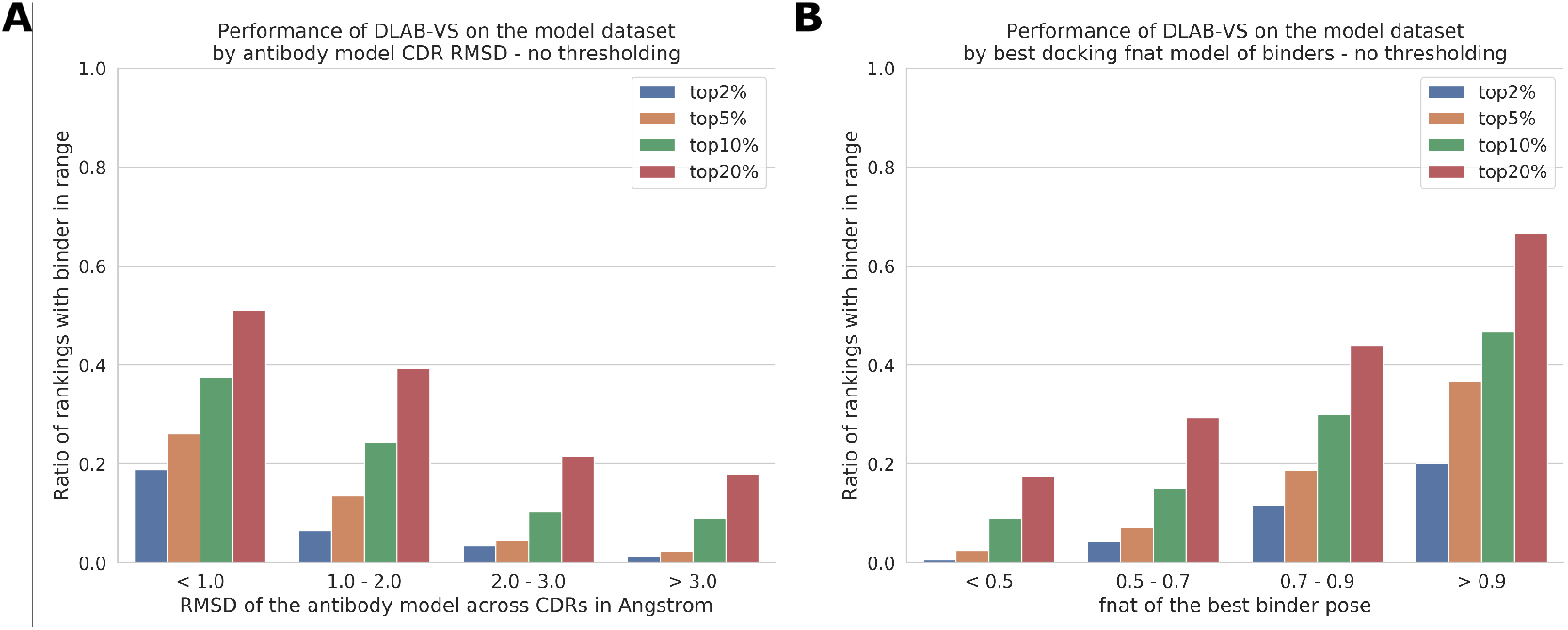
Dependence of the performance of DLAB-VS+ZDock on antibody model quality (A), measured via RMSD of the CDR regions to the corresponding crystal structure, and docking quality (B), measured via the highest fnat achieved in the top500 docking poses generated by ZDock. Both good antibody models and high quality docking poses correlate with high classification performance.

**Supplementary Figure 12.**
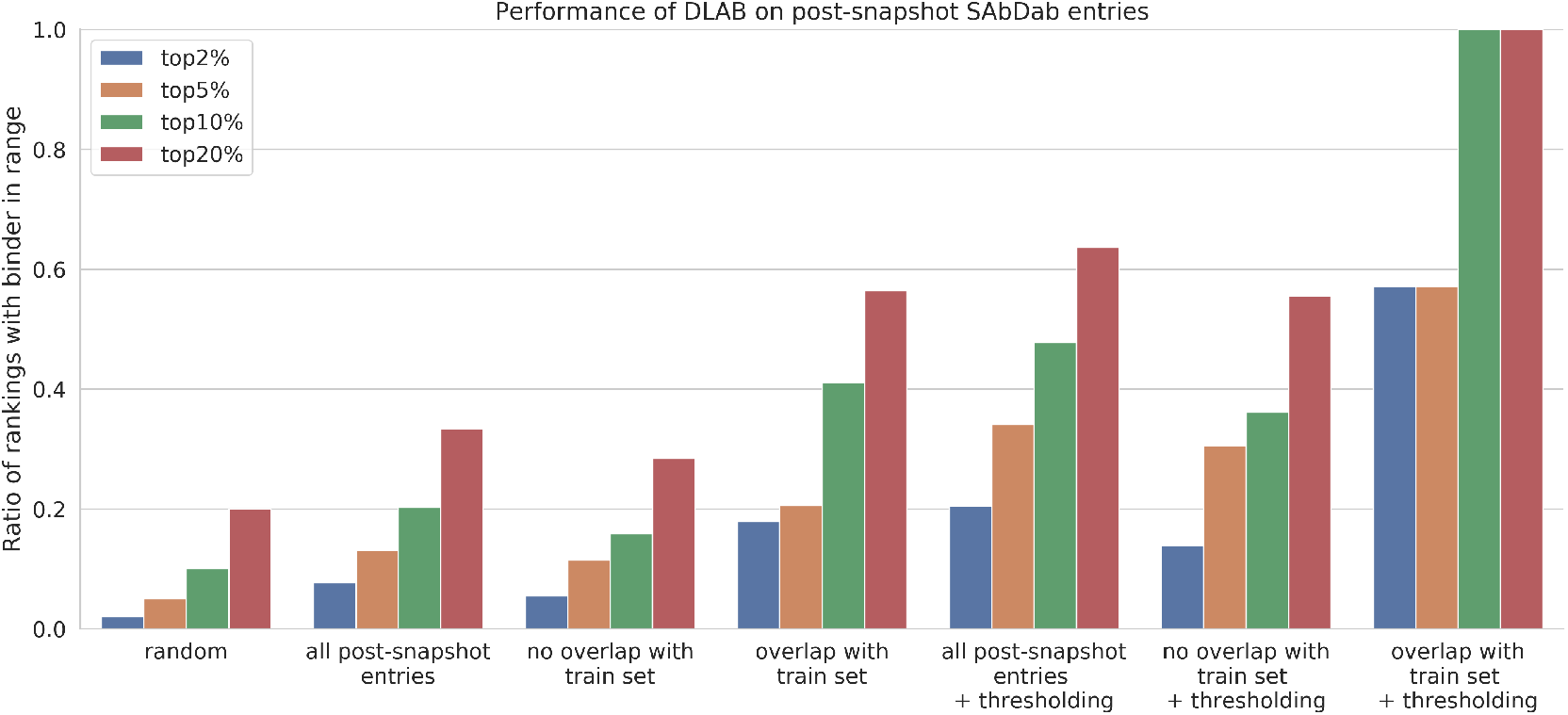
Full overview of the performance of DLAB-VS+ZDock on the post-snapshot model data set. As in the main text figure, the bars show the fraction of antigens within the data set for which the correct binder is ranked by DLAB-VS within the top 2%, top 5%, top 10% or top 20% respectively. We compare the random expectation value (“random”) to the whole post-snapshot model set (“all post-snapshot entries”), the antigen targets within the set for which the binding antibody does (“overlap with train set”) or does not (“no overlap with train set”) cluster with at least one antibody from the model data set at 90% CDR sequence identity. For each of these three options, the improved performance upon using the DLAB-Re-max score to discard 80% of antigen targets as described above is shown as well (“+ thresholding”).

**Supplementary Figure 13.**
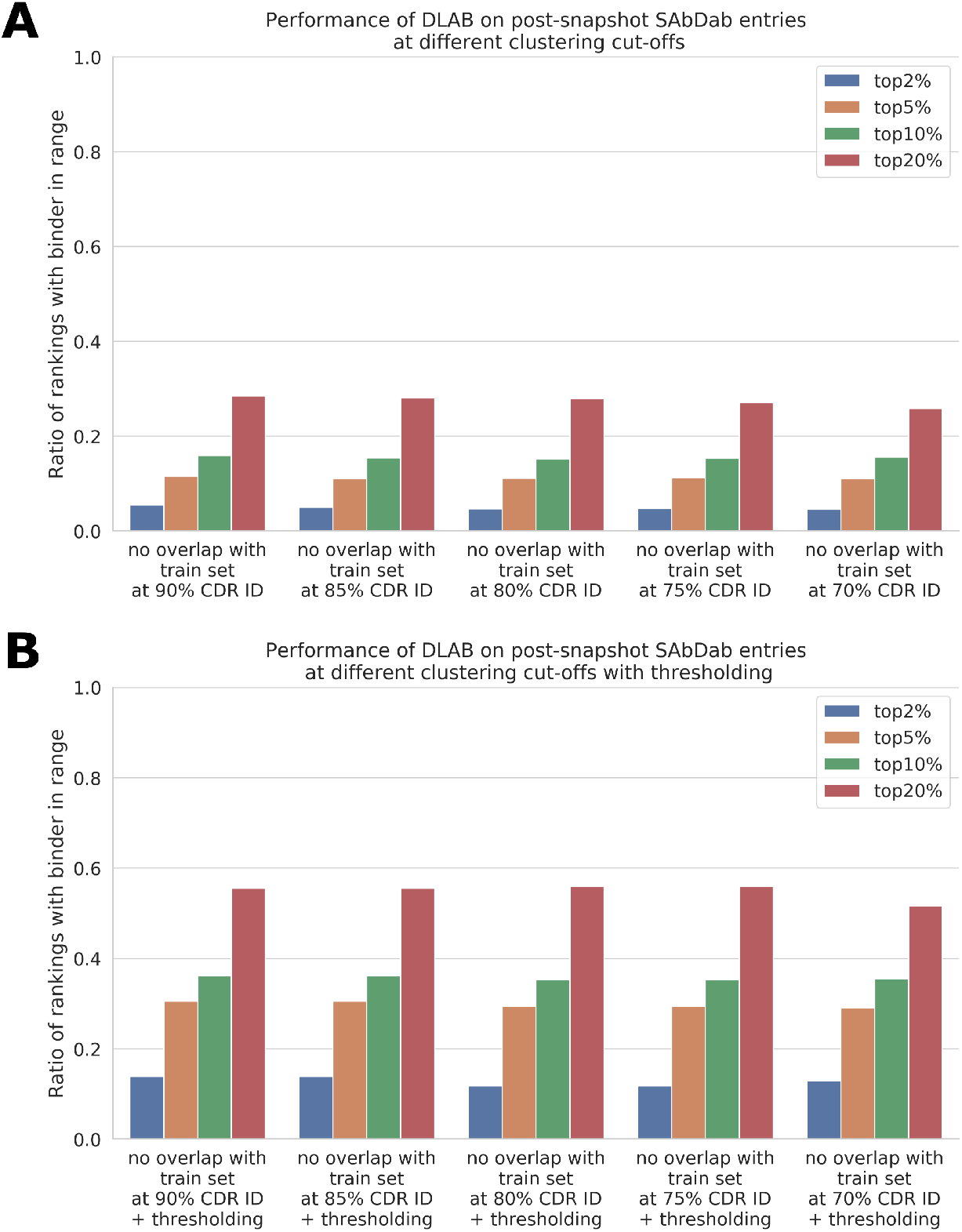
Performance of DLAB-VS+ZDOCK on the post-snapshot dataset at different percentage CDR sequence identity cutoffs for overlap with the training set without (A) and with (B) DLAB-Re thresholding. The performance of the DLAB-VS+ZDOCK model only marginally declines when limiting the allowed overlap to training set antibodies from 90% CDR ID to 70% CDR sequence identity, demonstrating that the model performance is generalisable.

